# Recent emergence of cephalosporin resistant *Salmonella* Typhi in India due to the endemic clone acquiring IncFIB(K) plasmid encoding *bla*_CTX-M-15_ gene

**DOI:** 10.1101/2023.07.05.547856

**Authors:** Tharani Priya T, Jobin John Jacob, V Aravind, Monisha Priya T, Bhavini Shah, Veena Iyer, Geeti Maheshwari, Urmi Trivedi, Anand Shah, Pooja Patel, Anushree Gaigawale, M Yesudoss, Pavithra Sathya Narayanan, Ankur Mutreja, Megan Carey, Jacob John, Gagandeep Kang, Balaji Veeraraghavan

**Affiliations:** Christian Medical College, Vellore, Tamil Nadu, India; Neuberg Supratech Reference Laboratories, Ahmedabad, Gujarat, India; Indian Institute of Public Health, Gandhinagar, Gujarat, India; Toprani Advanced Lab Systems, Vadodara, Gujarat, India; Unipath Speciality Laboratory, Akota, Vadodara, Gujarat, India; Zydus Hospitals, Ahmedabad, Gujarat, India; Pathocare Pathology Laboratory, Vadodara, Gujarat, India; Suburban Diagnostics India Pvt. Ltd, Andheri (w), Mumbai, India; Cambridge Institute of Therapeutic Immunology and Infectious Disease, Jeffrey Cheah Biomedical Centre, Cambridge, UK; Department of Infection Biology, Faculty of Infectious and Tropical Diseases, London School of Hygiene & Tropical Medicine, London, UK

**Keywords:** Ceftriaxone-resistant, *Salmonella* Typhi, Genome analysis, Plasmid, Antimicrobial resistance

## Abstract

The emergence and spread of *Salmonella* Typhi (*S*. Typhi) resistant to third-generation cephalosporins is a serious global health concern. In this study, we genomically characterized 142 cephalosporin-resistant *S*. Typhi strains isolated from India. Comparative genome analysis revealed the emergence of a new clone of ceftriaxone-resistant *S*. Typhi harboring three plasmids of the incompatibility groups IncFIB(K), IncX1, and IncFIB(pHCM2). Among these, the IncFIB(K) plasmid confers resistance to third-generation cephalosporins through the *bla*_CTX-M-15_ gene, along with other resistance determinants such as aph(3“), aph(6’), *sul2*, *dfrA14, qnrS,* and *tetA*. Phylogenetic analysis showed that the isolates from Gujarat (n=140/142) belong to a distinct subclade (genotype 4.3.1.2.2) within genotype 4.3.1.2 (H58 lineage II). SNP-based phylogenetic analysis of the core genes in IncFIB(K) reveals a close genetic relationship between the plasmid backbone and that of IncFIB(K) from other Enterobacterales, suggesting that the H58 lineage II possesses the capability to acquire MDR plasmids from these organisms. This could indicate the potential onset of a new wave of ceftriaxone-resistant *S.* Typhi in India. The implementation of control measures such as vaccination, and improved water, sanitation, and hygiene (WASH) systems, is crucial in areas where MDR or XDR *S*. Typhi strains are prevalent to curb the spread and impact of these resistant strains.

**Importance:** Typhoid fever remains a significant public health problem in many parts of the world, particularly in resource-limited settings with inadequate sanitation and limited access to clean water. The emergence of MDR *S. Typhi* resistant to first-line antibiotics (chloramphenicol, ampicillin, and cotrimoxazole) and fluoroquinolones marked the initial treatment challenge, leading to reliance on azithromycin and third-generation cephalosporins. Recently, researchers identified the emergence of XDR typhoid in Pakistan that is resistant to five classes of antibiotics including third-generation cephalosporins. In this study, we report a recent emergence of *S*. Typhi isolates resistant to third-generation cephalosporins. Genomic characterization and evolutionary analysis of the new strains revealed a recent acquisition of *blaCTX-M* carrying IncFIB(K) plasmid led to the emergence of ceftriaxone-resistant *S.* Typhi. Due to the ongoing nature of this outbreak, the data from this study deserves further consideration in order to control its spread in India.

## Introduction

Enteric fever is a systemic febrile illness caused by the human-restricted pathogens *Salmonella enterica* serovar Typhi and Paratyphi A, B, and C (Crump and Mintz, 2010). It continues to be a major cause of morbidity and mortality in low-and middle-income countries in South Asia, Southeast Asia, and Africa (Antillón et al., 2017). In most cases, typhoid infection is contracted from contaminated food or water, rather than through direct contact with an infected person. Globally, typhoid fever is estimated to cause approximately 11 to 20 million cases and results in between 28,000 and 161,000 deaths annually (WHO, 2023; Mogasale et al., 2014). Another estimate from the Institute for Health Metrics and Evaluation (IHME) in 2019 suggested a global burden of 9.24 million cases of typhoid fever, associated with around 110,000 deaths (https://www.healthdata.org/results/gbd_summaries/2019/typhoid-fever-level-4-cause). Notably, region-wise data indicates that high-burden countries in South Asia account for roughly 70% of the global burden of enteric fever (Stanaway et al., 2019).

Historically, antimicrobial therapy was the most effective method of combating typhoid; however, this has been threatened by the emergence of antimicrobial-resistant (AMR) strains (Browne et al., 2020). During the late 1970s and early 1980s, multidrug-resistant (MDR) strains of *S*. Typhi became a major global concern (Marchello et al., 2020). Consequently, fluoroquinolones became the drugs of choice in the treatment of MDR *S*. Typhi. However, the emergence of fluoroquinolone non-susceptible *S*. Typhi has shifted the treatment options to azithromycin and third-generation cephalosporins (3GC) (Britto et al., 2018; Parry et al., 2019). These two drugs have continued as the ‘last resort’ antibiotics for treating enteric fever in the endemic regions of South Asia. A recent outbreak of extensively drug-resistant (XDR) *S*. Typhi strains (resistant to three first-line antibiotics, quinolones, and 3GCs) in Pakistan has caused treatment recommendations to shift in favor of azithromycin (Klemm et al., 2018).

The introduction of genome-based surveillance for *S*. Typhi based on GenoTyphi phylogenetic genotyping scheme has greatly improved our knowledge of the origins, transmission, and antibiotic resistance profiles of *S*. Typhi isolates (Wong et al., 2016). The current version of GenoTyphi scheme (Dyson and Holt, 2021) classifies the *S*. Typhi population into four major lineages, which are further divided into over 75 distinct clades and subclades. According to the population structure of *S*. Typhi strains, there are 85 haplotypes (H1 to H85), with haplotype 58 (genotype 4.3.1) being particularly notable for its high pathogenicity and multidrug resistance (Wong et al., 2015; 2016). During the past few decades, H58 *S*. Typhi has become the globally dominant lineage due to its efficient transmission and persistence within human populations (Pragasam et al., 2020; Carey et al., 2022). The intercontinental spread of MDR *S*. Typhi (genotype 4.3.1.1 or H58 lineage 1) was largely driven by the acquisition of an incompatibility (Inc)HI1 plasmid carrying multiple antimicrobial resistance determinants (*bla*_TEM_, *cat*, *dfrA*, *sul2*) (Holt et al., 2011; da Silva et al., 2022). Similarly, with the introduction of ciprofloxacin as the primary treatment for typhoid, a distinct sublineage (genotype 4.3.1.2 or H58 lineage 2) has emerged, characterized by fluoroquinolone non-susceptibility (FQNS) and the presence of double/triple-point mutations in the quinolone resistance determining regions (QRDRs) (Pham et al., 2016). Further, the emergence of a new sub-lineage P1 (4.3.1.1.P1) of *S*. Typhi in Pakistan has raised concerns due to its resistance to third-generation cephalosporins in addition to first-line antimicrobials and fluoroquinolones. Extensively drug-resistant (XDR) isolates belonging to this sub-lineage carry *bla*_CTX-M-15_, which is mobilized by IncY plasmids (Klemm et al., 2018). Complicating matters further, the development of point mutations in the *acrB* gene (R717Q/L) has given rise to azithromycin-resistant *S*. Typhi, posing a significant threat to the last available oral therapy option (Hooda et al., 2019; Duy et al., 2020).

Genomic epidemiology studies of *S*. Typhi have demonstrated the evolution and spread of multiple distinct lineages on a regional and global scale (Dyson and Holt, 2021). The regional dominance of H58 sublineages such as 4.3.1.1 (MDR) and 4.3.1.1P1 (XDR) in Pakistan, 4.3.1.2 (FQNS strains) in India and Nepal, and IncFIB(K) carrying genotype 4.3.1.3 in Bangladesh are well documented (Carey et al., 2023). Contrary to this, the population structure of *S*. Typhi in Africa shows H58 in eastern and southern Africa, while west and central Africa consist primarily of non-H58 lineages or distinct genotypes of H58 (Park et al., 2018; Carey et al., 2023). While the XDR lineage (4.3.1.1P1) has become dominant in Pakistan (70% in 2020), it has not yet become established elsewhere. Conversely, ceftriaxone-resistant (CefR) *S*. Typhi strains reported from India primarily belong to sub-clade 4.3.1.2 (recently renamed as 4.3.1.2.1.1) and carry *bla*_SHV-12_ mobilized by the IncX3 plasmid backbone (Jacob et al., 2021, Argimón et al., 2022). Over the past few years, ESBL-harboring plasmids belonging to different incompatibility types have been observed in *S*. Typhi isolates in India. However, their role in the evolution of antimicrobial resistance remains poorly understood (Jacob et al., 2021). In this study, we aimed to analyze the genomic epidemiology and evolutionary origin of 142 cases of ceftriaxone-resistant *S*. Typhi in India. We also explored recent plasmid transfer events and gene flow that led to the emergence of IncFIB(K) carrying cephalosporin-resistant *S*. Typhi in India.

## Methodology

### Ethical Statement

This study was approved by the Institutional Review Board (IRB) of Christian Medical College, Vellore (IRB Min No: 13489 dated 28.10.2020).

### Study setting and design

The Surveillance of Enteric Fever in India (SEFI) study, which concluded its first phase from 2018 to 2020, monitored the changes in the antimicrobial susceptibility of *S*. Typhi and *S*. Paratyphi A in India. To continue tracking antimicrobial resistance patterns, we initiated a smaller-scale second phase of the SEFI network’s surveillance in 8 cities, commencing in March 2022. In Ahmedabad, Gujarat, Neuberg Supratech Reference Laboratories (NSRL), in collaboration with the Indian Institute of Public Health, Gandhinagar, functioned as a reference laboratory for hospitals, clinics, and diagnostic centers in the region, undertaking the culturing and preliminary identification of isolates. Initial cases of ceftriaxone-resistant *S*. Typhi were reported from NSRL and confirmed at the Department of Clinical Microbiology, Christian Medical College, Vellore. After issuing an alert to clinicians and laboratories about the new strain, isolates from other parts of the state (Vadodara) were largely identified by Toprani Advanced Lab Systems (n=86) and Unipath Specialty Laboratory (n=31). Once confirmed as *S*. Typhi by conventional biochemical tests, samples were shipped to the central laboratory at the Department of Clinical Microbiology, Christian Medical College, Vellore, for confirmation and further testing. The present study examined a collection of 142 ceftriaxone-resistant *S*. Typhi isolates obtained from five clinical diagnostic laboratories located in Ahmedabad and Vadodara, India, between June 2022 and September 2023 (Suppl Fig. 1).

### Phenotypic characterization

Identification of *S*. Typhi isolates transferred to the central laboratory was re-confirmed using conventional biochemical tests, and serotyping was conducted using commercial anti-sera (BD Difco, USA) based on the Kauffmann-White scheme according to the manufacturer’s instructions. Phenotypic antimicrobial susceptibility testing was performed using the Kirby-Bauer disc diffusion method, and the zone diameter was measured and interpreted based on Clinical Laboratory Standards Institute (CLSI, 2023) guidelines. The tested antimicrobial agents included ampicillin (10 µg), chloramphenicol (30 µg), trimethoprim/sulfamethoxazole (1.25/23.75 µg), ciprofloxacin (5 µg), pefloxacin (5 µg), ceftriaxone (30 µg), cefixime (5 µg), and azithromycin (15 µg). In addition, the minimum inhibitory concentration (MIC) for ceftriaxone and azithromycin was determined using broth microdilution (BMD) and the results were interpreted using CLSI guidelines (CLSI, 2023).

### DNA extraction and whole genome sequencing (WGS)

Genomic DNA was extracted using the QIAamp® Mini Kit (QIAGEN, Hilden, Germany) following the manufacturer’s instructions. The purity and concentration of the extracted DNA were measured using Nanodrop One (Thermo Fisher, Waltham, USA) and Qubit Fluorometer using the dsDNA HS Assay Kit (Life Technologies, Carlsbad, USA). For short-read sequencing, genomic DNA was fragmented and the paired-end library was prepared using Illumina Nextera DNA Flex Library Kit and Nextera DNA CD Indexes (Illumina, Massachusetts, MA, USA). The libraries were pooled at equal molar concentration and sequenced on the Illumina Novaseq 6000 platform, yielding 2×150 bp paired-end reads (Illumina, San Diego, CA, USA). DNA tagmentation, library amplification, and clean-up were performed according to the manufacturer’s protocol. A subset of five isolates was subjected to sequencing using Oxford Nanopore Technology (ONT). The libraries for sequencing were prepared using the Ligation Sequencing Kit (SQK-LSK114) (ONT, UK) and sequenced on an ONT MinION device using the R10.4.1 flow cell and Q20 chemistry according to the manufacturer’s instructions. Duplex ONT read pairs were prepared for basecalling using ONT’s duplex tools (https://github.com/nanoporetech/duplex-tools) and raw nanopore reads were basecalled with Guppy (https://pypi.org/project/ont-pyguppy-client-lib/).

### QC, Assembly

Short reads generated from the Illumina platform were checked for quality using FastQC v0.12.1, and adapters and indexes were trimmed using Trimmomatic, v0.39. Contamination detection and filtering were performed by Kraken v1.1.1 (https://github.com/DerrickWood/kraken) followed by sequence coverage. The high-quality reads (Phred score >30) were assembled using SPAdes (https://github.com/ablab/spades) with option--careful. Long-read data generated from the ONT sequencer were subsampled to an average coverage depth of 100× using Rasusa v0.7.1. The reads were assembled using the Hybracter pipeline (v0.8.0; Bouras et al., 2024), which improved ONT read quality using Filtlong. The pipeline performed assembly of the long reads with Flye, followed by polishing with Medaka. This was further improved by polishing with Illumina short reads using the Polypolish and PyPolca careful modules (https://github.com/gbouras13/hybracter). The resulting complete genome assemblies were assessed by QUAST v5.2.0 and seqkit v2.4.0. Default parameters were used unless otherwise mentioned.

### Comparative genomics

From the global collection of *S.* Typhi genomes (n=415), reads representing all major genotypes were obtained from the European Nucleotide Archive (ENA; http://www.ebi.ac.uk/ena) or the Sequence Read Archive (SRA) based on the curated data provided by the Global Typhoid Genomics Consortium (https://bridges.monash.edu/articles/dataset/Global_Typhoid_Genomics_Consortium_2022_-_Genome_Assemblies/21431883) (Carey ME et al., 2023). Genomes were classified according to their sequence types using the Multilocus Sequence Typing (MLST) pipeline offered by the Center for Genomic Epidemiology (https://cge.cbs.dtu.dk/services/MLST/). *S.* Typhi genotypes were determined using the GenoTyphi scheme (https://github.com/katholt/genotyphi) (Wong et al., 2016). Antimicrobial resistance genes were identified from the genome sequences using the BLASTn-based ABRicate v0.8.10 program (https://github.com/tseemann/abricate) with the AMRFinderPlus database v3.11.14 (Feldgarden et al., 2019). Plasmids were identified and typed using PlasmidFinder (https://cge.cbs.dtu.dk/services/PlasmidFinder/) and MOB-suite pipelines (https://github.com/phac-nml/mob-suite) (Carattoli et al., 2014; Robertson J et al., 2020). The complete list of raw sequence data is available in the Suppl Table 1.

### Phylogenetic analysis

The assembled contigs (*n=574*) were mapped to the reference genome of *S.* Typhi CT18 (GenBank: AL513382.1) using Snippy v4.6.0 (https://github.com/tseemann/snippy) (Seemann, 2015). The whole genome SNP alignment was filtered for recombination using Gubbins v3.3.3 (https://github.com/nickjcroucher/gubbins) (Croucher et al., 2015). Single nucleotide polymorphisms (SNPs) were then called using SNP-sites v2.5.1 (https://sanger-pathogens.github.io/snp-sites/) (Page et al., 2016). Subsequently, a maximum-likelihood phylogeny was inferred from the alignment using RAxML-NG v1.2.1 (https://github.com/amkozlov/raxml-ng) under the GTR+GAMMA model with 200 bootstraps. The resulting phylogenetic tree was visualized and annotated using the Interactive Tree of Life software (iTOL v.5) (Letunic and Bork, 2021).

### Plasmid characterization

We identified 288 plasmid sequences typed as IncFIB(K) from public databases including NCBI’s RefSeq database (https://www.ncbi.nlm.nih.gov/), the PLSDB database (Schmartz et al., 2022), and COMPASS reference database (Douarre et al., 2020). IncFIB(K) plasmids previously reported from *S.* Typhi were retrieved from the curated data provided by the Global Typhoid Genomics Consortium (Carey et al., 2023). Contigs belonging to plasmids were extracted from the draft assembly using PlasForest v1.4 as previously described (Gomi et al., 2021). Incomplete and redundant sequences were removed manually, and plasmids were reconfirmed as IncFIB(K) using abricate v1.01 with the PlasmidFinder database (Giménez et al., 2022). Plasmid taxonomic units (PTUs) were assigned by COPLA (Redondo-Salvo et al., 2021) and plaSquid v1.0.0 (Giménez et al., 2022). Near-identical plasmids were detected with Mash tree v1.2.0 and outliers/singletons were removed from the analysis. The plasmid sequences were annotated with Prokka (https://github.com/tseemann/prokka) using the default parameters (Seemann et al., 2014). Annotated assemblies in the GFF3 format were used as input for pan-genome analysis using Panaroo (https://github.com/gtonkinhill/panaroo) in its “Strict” mode with a core threshold of 80 (Tonkin-Hill et al., 2020). The core genome alignment generated was used to construct plasmid core gene phylogeny using IQ-Tree v2.2.2.6 with parameters-m GTR+F+I+G4 (Minh et al., 2020). We performed RhierBAPS to define the IncFIB(K) population structure, and the phylogenetic cluster carrying IncFIB(K) was further redefined to fully understand plasmid evolution (Tonkin-Hill et al., 2018). The gene presence or absence in each genome obtained was grouped according to the phylogenetic lineages using twilight scripts (https://github.com/ghoresh11/twilight) with default parameters (Horesh et al., 2021).

### Determining plasmid stability

To evaluate the stability of plasmids IncFIB(K) and IncY within *S*. Typhi strains, we performed the following experimental procedure. Initially, the strains were cultured overnight in LB-ampicillin broth. The overnight culture served as the inoculum for fresh LB broth at a 1% volume, achieving an approximate initial cell density of 5 × 10^7^ CFU/ml (Wu et al., 2010). The cultures were then subjected to serial passaging every 12 hours over six days in antibiotic-free LB broth, maintained at 37°C with shaking at 150 rpm. Following each subculture, the bacterial cultures were serially diluted and plated on LB agar (Fan et al., 2022). To determine plasmid loss during the subculturing process, approximately 100 colonies from each culture were replica plated onto both LB agar and LB-ampicillin agar. This procedure allowed us to compare the growth on antibiotic-containing media against antibiotic-free media to assess plasmid retention. The plasmid stability test was conducted in triplicate for each strain, and the average number of colonies on the three LB plates was calculated to determine the retention rate of the plasmids.

## Data availability

Raw read data were deposited in the European Nucleotide Archive (ENA) under project accession number: PRJEB62461

## Results

### Phenotypic characterization

The ceftriaxone resistant *S.* Typhi isolates *(n=142)* investigated in this study were collected from febrile patients attending private healthcare facilities in Ahmedabad and Vadodara, India. Among the study cohort, 29 isolates were obtained from diagnostic laboratories in Ahmedabad, 113 isolates were collected from Vadodara, while one isolate was obtained from a diagnostic facility in Mumbai (Suppl Fig 1). Antimicrobial susceptibility profiling revealed that all except one (*n=141/142*) study isolates exhibited resistance to ampicillin, co-trimoxazole, ciprofloxacin, and ceftriaxone, while showing susceptibility to chloramphenicol and azithromycin. The isolate from Mumbai (*n=1*) was susceptible to co-trimoxazole but resistant to ceftriaxone. The minimum inhibitory concentration of ceftriaxone was determined and found to be >512 µg/ml.

### Genotype assigning

The genomes of 140/142 *S*. Typhi isolates were characterized by the GenoTyphi scheme (https://github.com/typhoidgenomics/genotyphi). All study isolates belonged to the prevalent subclade 4.3.1 genotype (H58), further specified as a unique genotype 4.3.1.2.2 based on the marker G3014988A. This novel genotype has been incorporated into the GenoTyphi scheme, enhancing its utility for early detection in future surveillance studies.

### Population structure of *S*. Typhi

The population structure of ceftriaxone resistant *S.* Typhi from the study collection was inferred from a core gene SNP-based phylogeny. The phylogenetic relationship of *S.* Typhi isolates in comparison to a curated set of global genome collection (*n=415*) and contextual isolates from Gujarat (*n=16*) highlighted the placement of isolates within clades corresponding to genotype 4.3.1 (H58) (Suppl. Fig. 2). Further analysis at the sublineage level revealed that majority of *S*. Typhi isolates (*n=140/142*) from Gujarat are closely related, forming a distinct subclade (4.3.1.2.2) within genotype 4.3.1.2 (H58 lineage II). The SNP analysis identified 16 core SNP differences between 4.3.1.2.2 the nearest neighbor (Suppl Table 2). The sole isolate obtained from Mumbai (1CTRMumbai), which was susceptible to co-trimoxazole, was assigned to the parent clade 4.3.1 while another (A99070) belonged to genotype 4.3.1.1 (Fig. 1).

**Fig 1:**
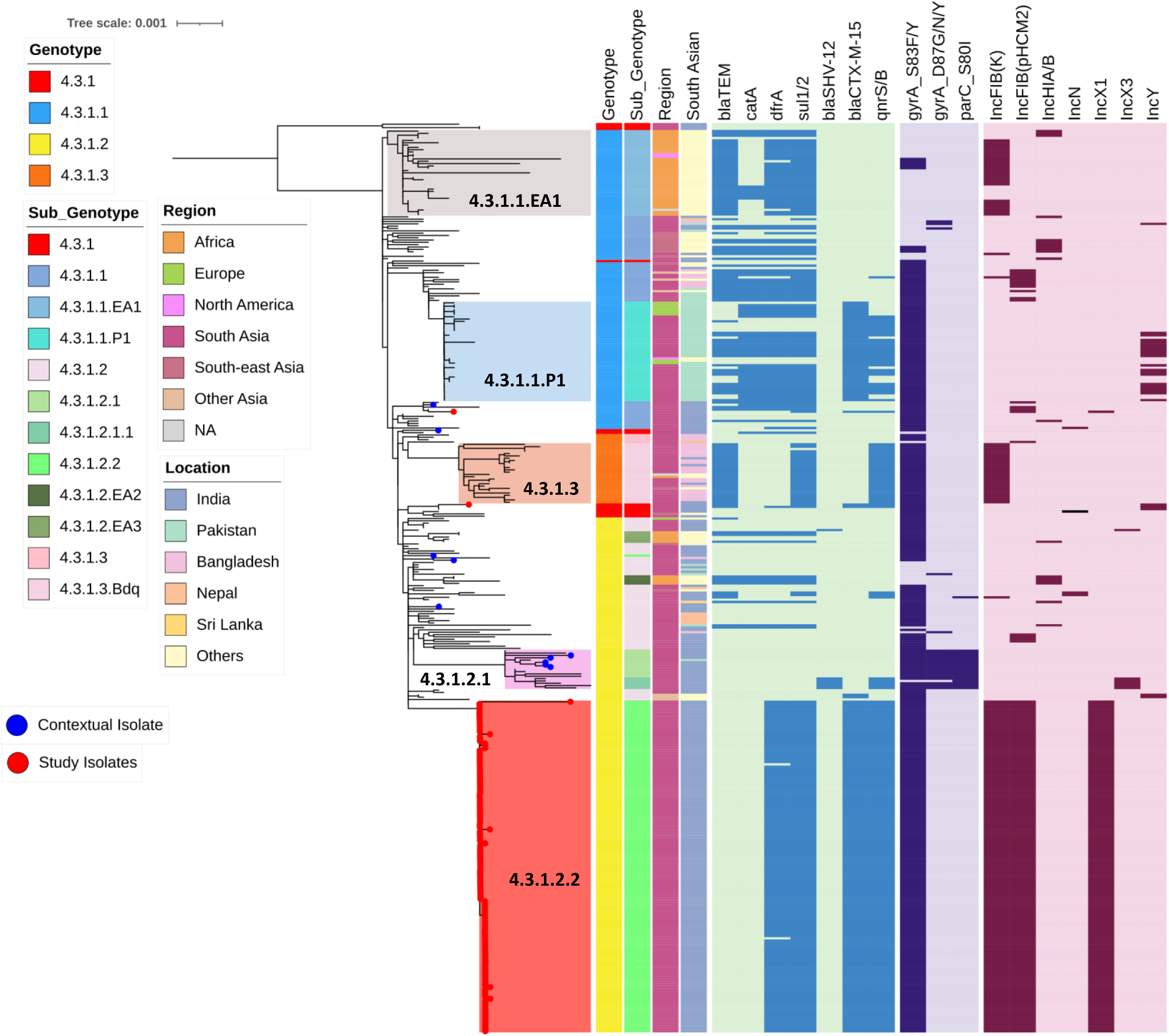
Population structure of H58 *S*. Typhi isolates. Maximum likelihood phylogenetic tree based on core genome sequence SNPs of 247 global H58 *S*. Typhi mapped to *S*. Typhi CT18 and rooted to outgroup isolates (Genotype 4.3.1). Major H58 sub-lineages such as 4.3.1.1EA1, 4.3.1.1P1, 4.3.1.3 and 4.3.1.2.1 are shaded in different colours. Red-colored dots at the tip of the branches indicates the position of this study isolates. Metadata are labeled as color strips and key for each variable were mentioned. Strip 1, 2 and 3 indicate the Genotype, Region of isolation and location in South Asia of each genomes. Heatmap represents the plasmid mediated resistance genes, QRDR mutations that confer resistance to fluoroquinolone and the presence of plasmids. The scale bar indicates substitutions per site. Color keys for all the variables are given in the inset legend. The tree was visualized and labeled using iTOL (https://itol.embl.de/).

Further, AMR gene analysis revealed that the study isolates carried *bla*_CTX-M-15_, *qnrS1*, *sul2*, *dfrA14*, *tet(A)* on an IncFIB(K) plasmid. Mutation analysis on the QRDR region showed single point mutation in *gyrA*: S83F conferring non-susceptibility to fluoroquinolones. This collection exhibits an accumulation of acquired AMR genes, in contrast to the most susceptible contemporaneous *S*. Typhi collections from India. No known mutations associated with azithromycin resistance were detected among any of the *S*. Typhi isolates

### Comparison of plasmids and AMR bearing region

A comparison of the complete circular IncFIB(K) plasmid from the study isolates (CP168964) with reference plasmids available in NCBI revealed the acquisition of antimicrobial resistance determinants within a broader genomic context (Fig. 2). Notably, while the reference IncFIB(K) plasmids from *E. coli* (CP116920) and *S. flexneri* (CP097833) carried the *bla*_TEM-1_ gene, it was absent in the study isolates. A detailed examination of the genetic environment surrounding the *bla*_CTX-M-15_ gene is provided in Suppl Fig 3. Although the genetic structures carrying *bla*_CTX-M-15_ in IncFIB(K) and IncY plasmids are similar, there are minor variations in their genetic arrangement. In the IncFIB(K) plasmid, the mobile element located upstream of *bla*_CTX-M-15_ is the IS1380 family ISEcp1, whereas, in the IncY plasmid, it is the IS1380 family ISEc9 transposase. Downstream of *bla*_CTX-M-15_, both plasmids contain the mobile elements wbuC, Tn3 family transposase, *qnrS1*, and an ISKra4-like element ISKpn19 family transposase. Notably, the IS1380 family appears to play a crucial role in the transfer of *bla*_CTX-M-15_ between plasmids within Enterobacteriaceae. Additionally, a comparison of IncFIB(K) and IncY plasmid sequences, covering replication mechanisms, stability, conjugation abilities, and AMR-bearing regions, is presented (Suppl Table 3).

**Fig 2:**
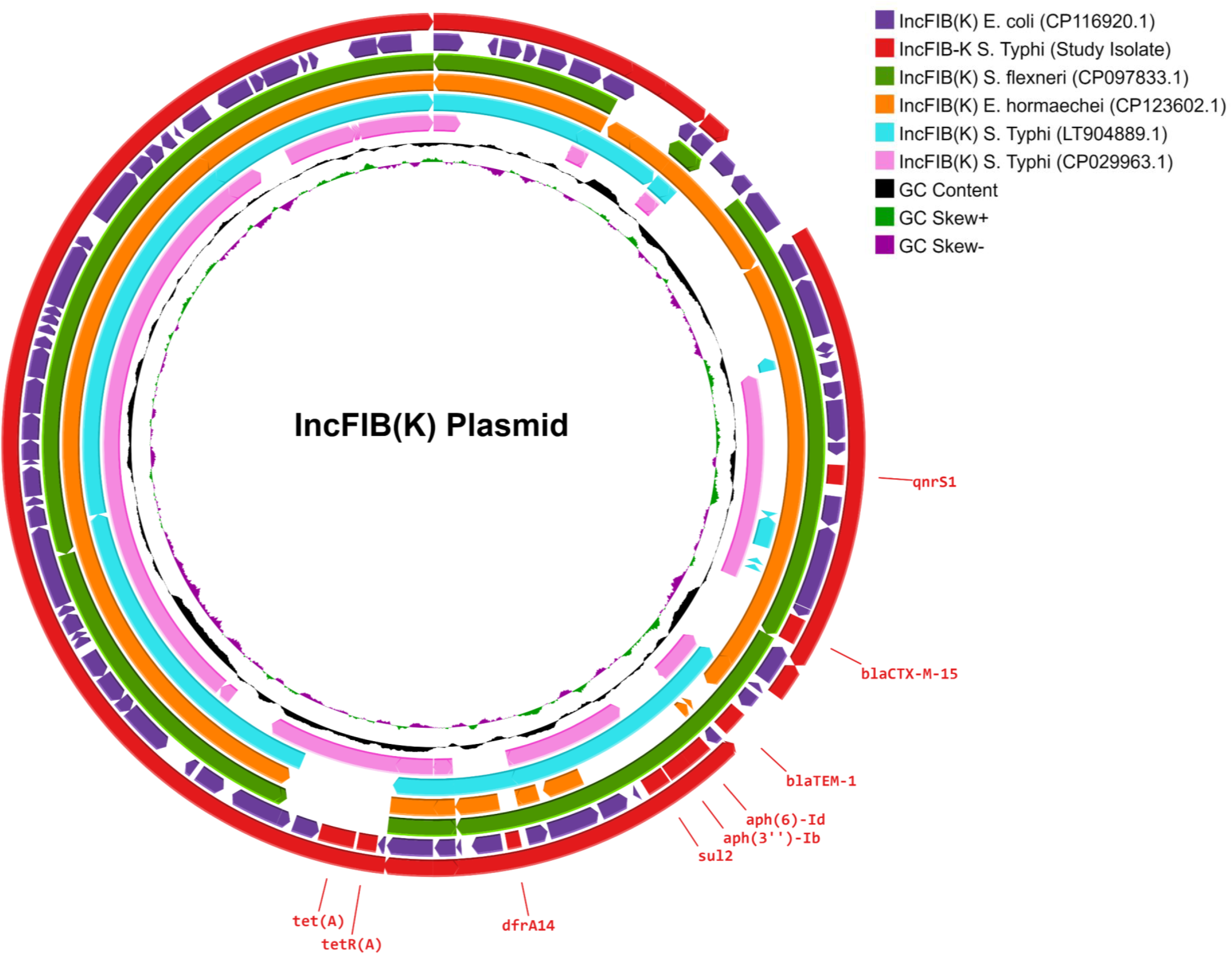
Comparison of IncFIB(K) plasmid. IncFIB(K) plasmid from ceftriaxone resistant *S*. Typhi isolates from Ahmedabad (Accession number: CP168964) is compared to plasmid of same incompatibility group from other Enterobacteriales [Accession number: CP116920(2023), CP097833(2022), CP123602(2023), LT904889(2017)] using Proksee (https://proksee.ca/). GC-content and GC-skew of the reference are depicted in the two innermost circles.

### Population structure of IncFIB(K) plasmids

Our curated dataset consists of IncFIB(K) plasmids (*n=276*) representing major Enterobacterales genera. The dataset includes plasmids from a diverse of bacterial hosts, with 53.9% from *K. pneumoniae*, 19.9% from *S*. Typhi, 9.6% from *E. coli* and 7.9% from other *Klebsiella* spp. Among the selected plasmids that are assigned as IncFIB(K) replicon types, MOB typing was performed and successfully classified into three MOB types, of which MOBV was the dominant MOB type. Notably twelve plasmids were assigned to multiple MOB types.

Pangenome analysis IncFIB(K) plasmid sequence showed a well-conserved backbone consisting of five genes (*repB*, *sopA*, *umuC/Y-family DNA polymerase*, *umuD* and *spo0J*) which formed a typical core gene set. After excluding outliers and incorporating our five newly sequenced IncFIB(K) plasmids, along with IncFIB(K) plasmids from other *S*. Typhi isolates, a final set of n=288 plasmid sequences were used for constructing a plasmid core gene phylogeny (Suppl Table 4). The genetic relatedness of selected IncFIB(K) plasmids revealed a diverse population structure with six major (level 1) BAPS clusters (Fig. 3a). Within the plasmid phylogeny, IncFIB(K) plasmids identified in the study isolates (Accession Number: CP168964) were positioned within cluster 1. This cluster is characterized by multiple bacterial hosts and is distinguished by a separation of four core SNPs (Suppl Table 5). To identify the near identical IncFIB(K) plasmids that were similar to those reported from study isolates, the entire plasmid sequences within BAPS cluster 1 (*n=76*) were examined. Within this cluster, 50 isolates formed a distinct sub-cluster, differentiated by an additional two core SNPs (Suppl Table 5) Core SNP distance-based plasmid phylogeny (Fig. 3b) demonstrates that plasmids from *E. coli* (OP242262), *S. flexneri* (CP097833) non-typhoidal Salmonella (NC_006856.1), and *Salmonella* Typhi (Tanzanian isolates) share a high degree of similarity with IncFIB(K) plasmids carried by the study isolates. Among the analyzed *S*. Typhi plasmids, LT904889 (a complete plasmid from a Tanzanian isolate) exhibits the closest genetic proximity.

**Fig 3:**
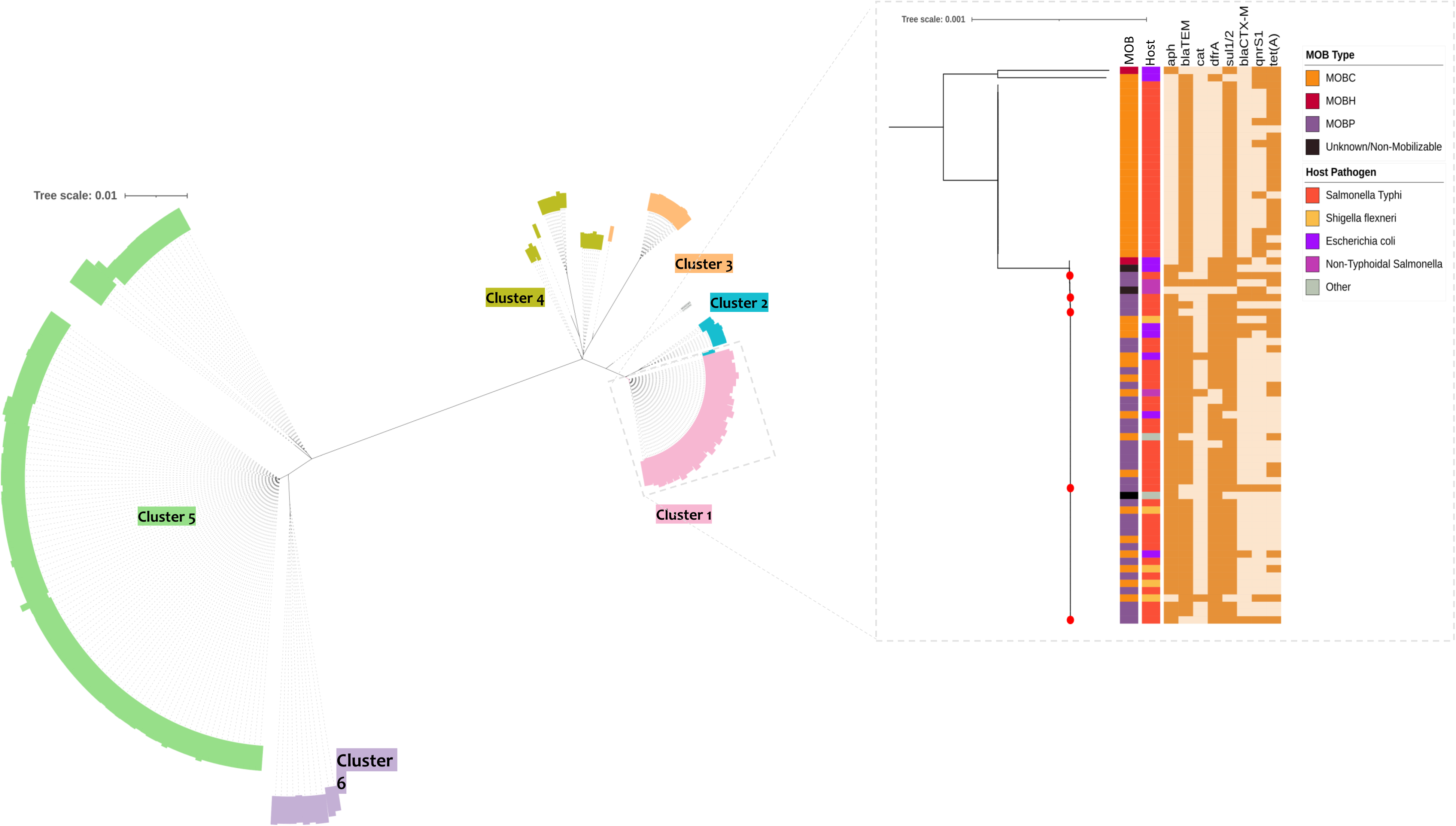
(a) An unrooted Maximum likelihood phylogenetic tree of global IncFIB(K) plasmids based on alignment of core genes. Phylogenetic clustering of different lineages is represented by RhierBAPS population clustering (level 1). Clusters are shaded in different colours. (b) A higher-resolution diagram of cluster 1 where IncFIB(K) plasmids from Gujarat were located. Metadata are labeled as color strips and key for each variable were mentioned. Strip 1 and 2 indicate the MOB type, and host organism. Heatmap represents the plasmid mediated resistance genes. Red-colored dots at the tip of the branches indicate the position of plasmids identified in this study.

### Plasmid stability analysis

The stability of the IncFIB(K) and IncY plasmids in their natural host, *S*. Typhi, was assessed through a series of experimental passage experiments over six days in antibiotic-free LB broth. The strain harboring the IncFIB(K) plasmid 154_NSRL(CP169080.1) exhibited high plasmid stability. After six days of serial passaging, 97% ± 1.5% of the colonies retained the plasmid, as evidenced by growth on LB-ampicillin agar. Similarly, the IncY plasmid also demonstrated high stability. After six days of serial passaging, 91% ± 3.2% of the strain carrying the IncY plasmid (CTR_R_Mumbai) retained the plasmid. These results indicate that both the IncFIB(K) and IncY plasmids are highly stable within the *S*. Typhi strains under the experimental conditions, with minimal plasmid loss observed over the duration of the study.

## Discussion

From June 2022 to September 2023, we identified 142 isolates of third-generation cephalosporin-resistant *S*. Typhi in India. The outbreak was initially detected as part of the SEFI phase 2 surveillance program for typhoid fever, which focuses on monitoring drug-resistant strains. Phenotypic analysis revealed resistance to ampicillin, trimethoprim-sulfamethoxazole, ciprofloxacin, and ceftriaxone, with susceptibility to chloramphenicol and azithromycin (Table 1). The resistance profile differs from classical extensively drug-resistant (XDR) strains, which are resistant to all first-line agents, fluoroquinolones and ceftriaxone (Klemm et al., 2018). Our study aimed to elucidate the population structure of the emerging *S*. Typhi clone and investigate the genetic background of plasmid transfer through whole genome sequencing (WGS) and comparative genome analysis.

Analysis of *S*. Typhi genome sequences revealed that the majority of ceftriaxone-resistant strains from Gujarat (*n=140/142*) belonged to genotype 4.3.1.2 and further subtyped to 4.3.1.2.2 (Fig. 1). The genotype 4.3.1.2 has been previously reported in multiple genome surveillance studies in India (Wong et al., 2015; Pragasam et al., 2020; Britto et al., 2020; Pragasam et al., 2021). However, genomic surveillance studies did not adequately represent circulating *S*. Typhi strains in western India, particularly in Ahmedabad or Vadodara. To address this gap, we sequenced 16 ceftriaxone-susceptible isolates (contextual) from Ahmedabad over the same period for comparative analysis. Five of these isolates were found to be closely related to the ceftriaxone-resistant strains, differing by only 26 SNPs (Suppl Fig: 4). This suggests that the endemic H58 lineage II clone circulating locally may have acquired an ESBL-encoding AMR plasmid. Meanwhile, the single isolate of ceftriaxone-resistant *S*. Typhi from Mumbai belonged to genotype 4.3.1, which has been previously detected in circulation in Northern India. (Sah et al., 2020; Dahiya et al., 2023).

One of the key observations was the plasmid profile of the emergent clone, which carries three plasmids that have not been previously reported in *S.* Typhi isolates. Among them, only IncFIB(K) carried resistance genes, while no resistance gene was found in IncFIB(pHCM2) and IncX1 plasmids. In particular, IncFIB(K) plasmid confers resistance to third-generation cephalosporins by means of *bla*_CTX-M-15_ gene, as well as other resistance determinants such as *aph(3“)*, *aph(6’*), *sul2, dfrA14*, *qnrS a*nd *tetA* (Fig. 2) Considering the restricted host range of *S*. Typhi, the acquisition of three plasmids was an unexpected event.

The evolutionary history of *S*. Typhi is characterized by genome degradation events, including deletions and inactivating mutations within coding sequences (McClelland et al., 2004; Holt et al., 2008; Wong et al., 2015). This trend has been corroborated by comparative genomic studies (Holt et al., 2009) and recent pan-genome analyses (Laing et al., 2017), which identify genome degradation, rather than gene acquisition, as the primary driver of *S*. Typhi evolution. Additionally, the genetic stability of *S*. Typhi is influenced by adaptive changes resulting in minimal genomic variation in the absence of antibiotic selection pressure (Jacob et al., 2021; Baddam et al., 2014). However, the emergence of resistant phenotypes has altered the evolutionary trajectory. MDR and XDR strains have acquired large plasmids such as IncHI1 and IncY (Wain et al., 2003; Klemm et al., 2018). Yet, plasmid-carrying clones have exhibited compensatory evolution, enabling the expansion of these clones over other phylogenetic lineages. (Wong et al., 2015). For example, IncHI1 plasmid acquisition drove the emergence of MDR *S*. Typhi (H58 lineage I/genotype 4.3.1.1) in South Asia (Holt et al., 2011; Carey et al., 2023). Subsequent chromosomal integration of the MDR composite transposon facilitated plasmid-free strains, stabilizing the MDR phenotype across lineages (Dyson et al., 2019; Carey et al., 2023). In Pakistan, the stabilized MDR genotype (4.3.1.1) evolved into an XDR clone (4.3.1.1.P1) via IncY plasmid acquisition harboring *bla*_CTX-M-15_ and *qnrS1* (Klemm et al., 2018). This suggests that H58 lineage I, with a chromosomal MDR transposon and a single QRDR mutation (S83F), can acquire and maintain plasmids carrying AMR determinants.

Interestingly, the expansion and regional dominance of H58 lineage II (genotype 4.3.1.2) in India has been driven by mutations in the *gyrA* and *parC* genes, which confer non-susceptibility to fluoroquinolones (Pragasam et al., 2020). This dominance of lineages with reduced susceptibility to fluoroquinolones is linked to the high fluoroquinolone exposure in the region (Dahiya et al., 2014). Notably, these QRDR double/triple mutant strains rapidly outcompeted other lineages due to the fitness advantage gained during evolution (Baker et al., 2013). As a result, H58 lineage II isolates are hypothesized to less likely acquire plasmids due to the initial cost associated with plasmid carriage (Jacob et al., 2021). Nevertheless, H58 lineage II isolates were previously reported to carry plasmids encoding cephalosporin resistance genes. Sporadic isolates of IncX3 and IncN plasmids carrying H58 lineage II isolates were reported between 2015 and 2019 (Jacob et al., 2021; Argimón et al., 2022). However, plasmid-free isolates likely outcompeted those bearing plasmids due to the associated fitness cost. Contrary to our expectations of rare plasmid carriage in H58 lineage II, the newly circulating clone of ceftriaxone-resistant isolates surprisingly carried three plasmids, one of which harbored resistance determinants We hypothesize that the emergence of ceftriaxone-resistant isolates from Gujarat, India could be the result of a recent event of acquisition of multiple plasmids from another Enterobacteriaceae donor. The SNP-based phylogeny of the core genes in IncFIB(K) plasmid backbone identified six major clusters within the collection (Fig. 2). Based on a closer examination of cluster 1, where IncFIB(K) plasmids of study isolates were located, we were able to identify sequences of plasmids from *S*. Typhi, *S*. *flexneri, E. coli* and other *Salmonella* sp. with a pairwise distances of 0 to 1 core SNPs (Fig. 2 & 3). In particular, the IncFIB(K) plasmid carried by study isolates is closely related to the plasmid identified in a previously sequenced Tanzanian strain of *S*. Typhi (LT904889.1) (Ingle et al., 2019). These findings suggest that H58 lineage II has the ability to acquire MDR plasmids from other Enterobacteriaceae, potentially facilitated by adaptive evolutionary mechanisms.

The emergence of new *S*. Typhi clones that are resistant to last-line antibiotics poses a serious problem for typhoid treatment and management. In view of the increased prevalence of resistance to third generation cephalosporins, azithromycin is the only oral treatment option, as per the current regimen. Though azithromycin and carbapenems remain effective for the treatment of uncomplicated and complicated typhoid fever, sporadic reports of resistance to these antibiotics have already been reported from the endemic areas (Octavia et al., 2021; Sajib et al., 2021; Carey et al., 2021; Ain et al., 2022). As a result, clinicians are faced with a challenging situation when choosing antibiotics for the treatment of typhoid fever. Considering the burden of typhoid fever, Indian NITAGs have advised the nationwide introduction of typhoid conjugate vaccine in June 2022. In order to mitigate the emergence of drug-resistant bacteria and the occurrence of untreatable typhoid cases, it is imperative that we take immediate action to widely implement Typhoid Conjugate Vaccines (TCVs), with a primary emphasis on introducing them in regions where antimicrobial resistance (AMR) is more prevalent.

## Conclusion

Comparative genome analysis of 142 ceftriaxone-resistant *S*. Typhi strains has elucidated the population structure and resistance plasmid dynamics in India. The acquisition of various mobile genetic elements and different genetic structures related to antibiotic resistance highlights the ongoing genomic remodeling in *S*. Typhi. Our findings provide additional evidence that H58 lineage II *S*. Typhi, previously thought to have a stable genome, has undergone rapid evolution through the acquisition of MDR plasmids. This evolution, likely driven by the selective pressure of third-generation cephalosporin treatment, may signal the onset of a new wave of ceftriaxone-resistant *S*. Typhi in India.

The emergence of these resistant strains underscores the urgent need for comprehensive public health interventions. The introduction of typhoid conjugate vaccines (TCVs), along with improvements in water, sanitation, and hygiene (WASH) systems, is crucial in reducing the selection pressure that drives the emergence of increasingly resistant *S*. Typhi strains. These measures, combined with ongoing surveillance and prudent antibiotic use, can help mitigate the spread and impact of drug-resistant typhoid fever.

## Acknowledgments

We thank Prof. Nicholas Grassly, Imperial College London, for assistance with study design and research proposal development. We acknowledge Dr. Duncan Steele, Ms. Megan Carey & Dr. Supriya Kumar, Bill & Melinda Gates Foundation for their technical support throughout the study on behalf of SEFI consortium. We thank all the lab members involved in SEFI reference lab activities, especially Ms. Agila Kumari P, Ms. Baby Abirami S, Mr. Ayyanraj N, Ms. Yamini Umashankar, Mr. Praveen Thilagan, CMC Vellore for their contributions to phenotypic testing and stock culture maintenance. We also thank all members of the SEFI consortium and the Wellcome Trust Research Laboratory, CMC Vellore, for their efforts in procuring clinical isolates and curating metadata. This study was approved by the Institutional Review Board (IRB) of Christian Medical College, Vellore (IRB Min No: 10393 dated 30.11.2016). Study participants were informed of the purpose and objectives of the study prior to the study. Written informed consent is obtained from every participant in the study before using their data for further research purposes.

