## Supplementary figures and images for "Recent emergence of cephalosporin resistant *Salmonella* Typhi in India due to the endemic clone acquiring IncFIB(K) plasmid encoding *bla*_CTX-M-15_ gene"

### supplementary Fig 1

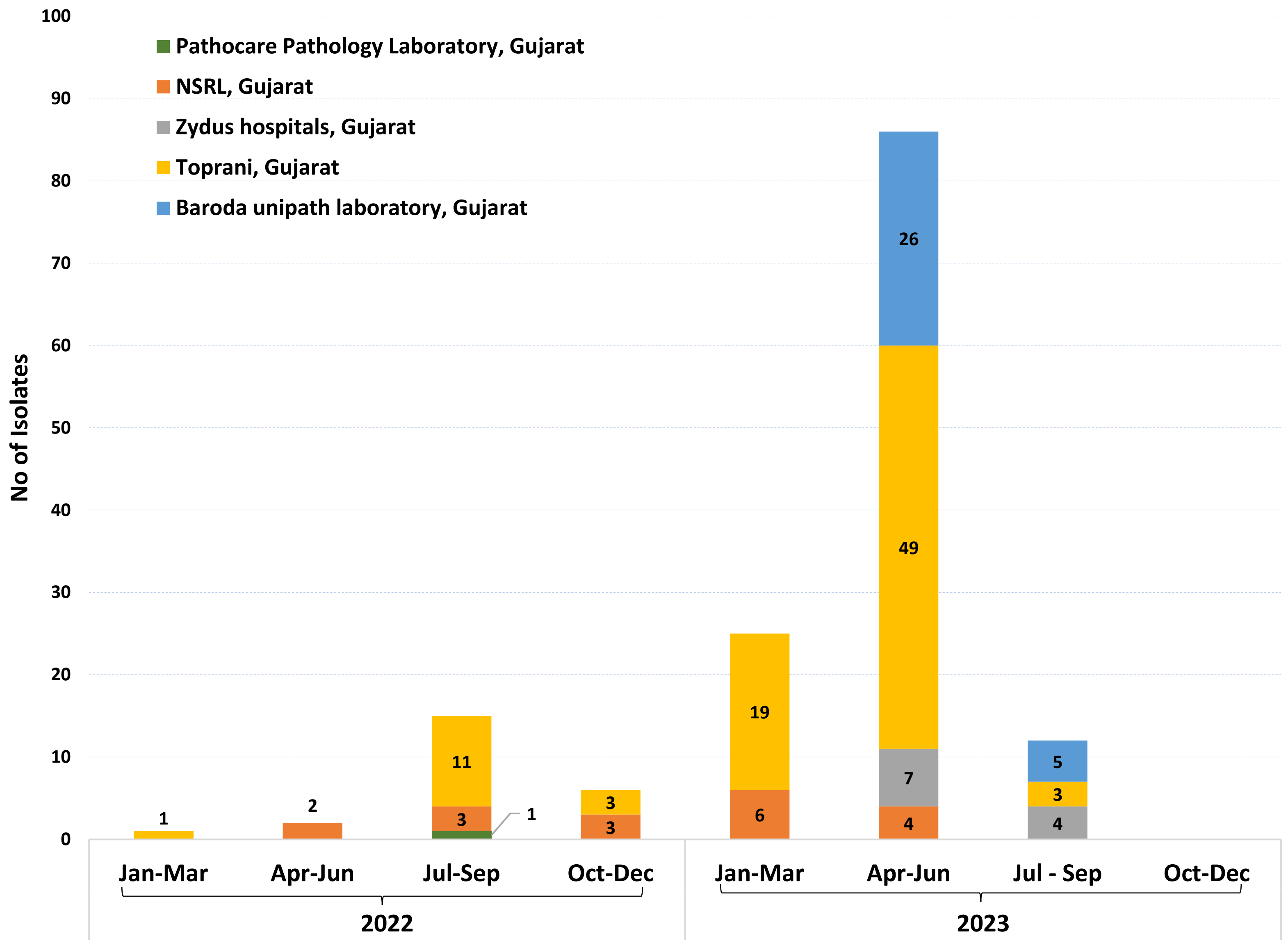

### supplementary Fig 2

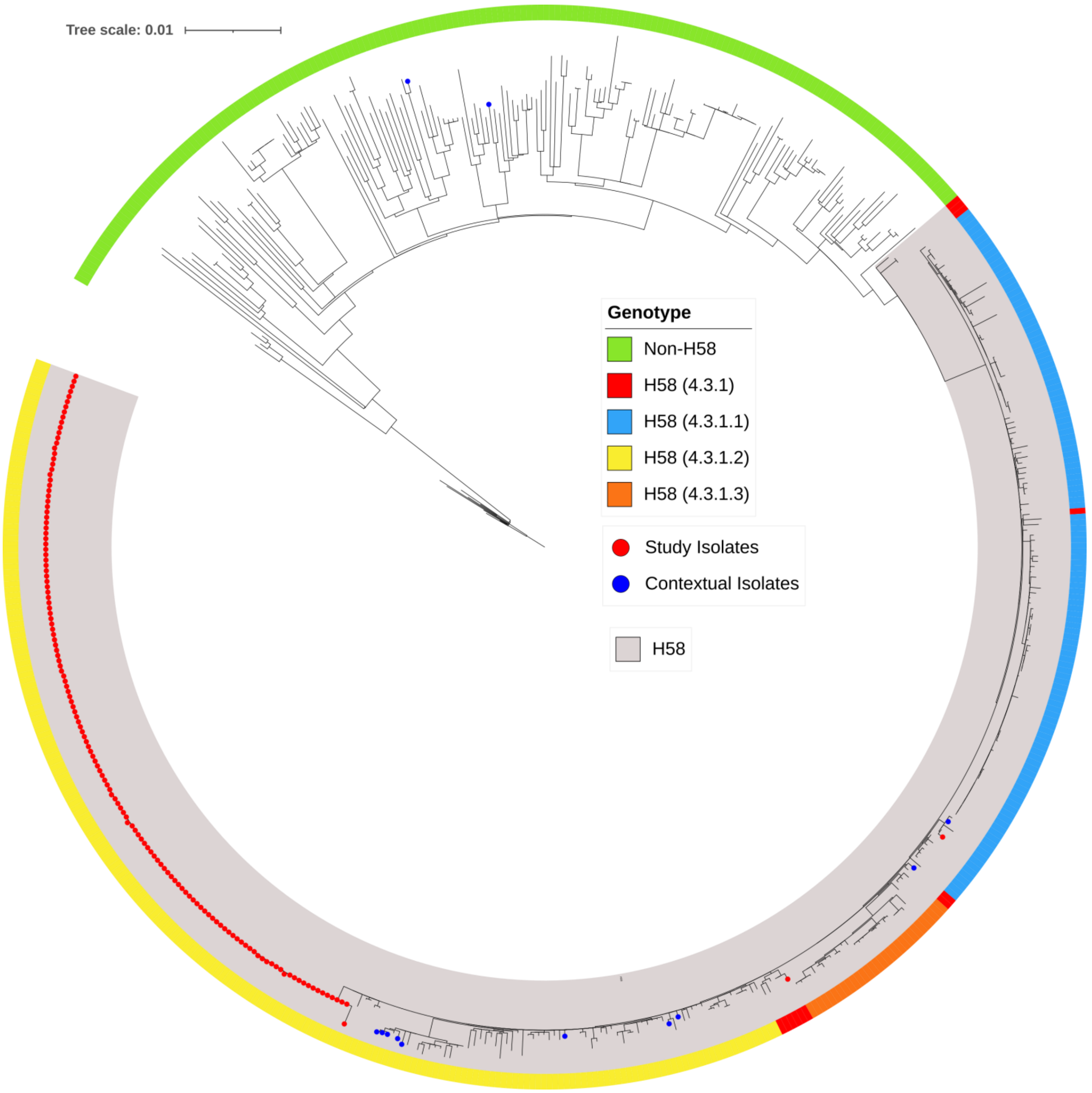

### supplementary Fig 3

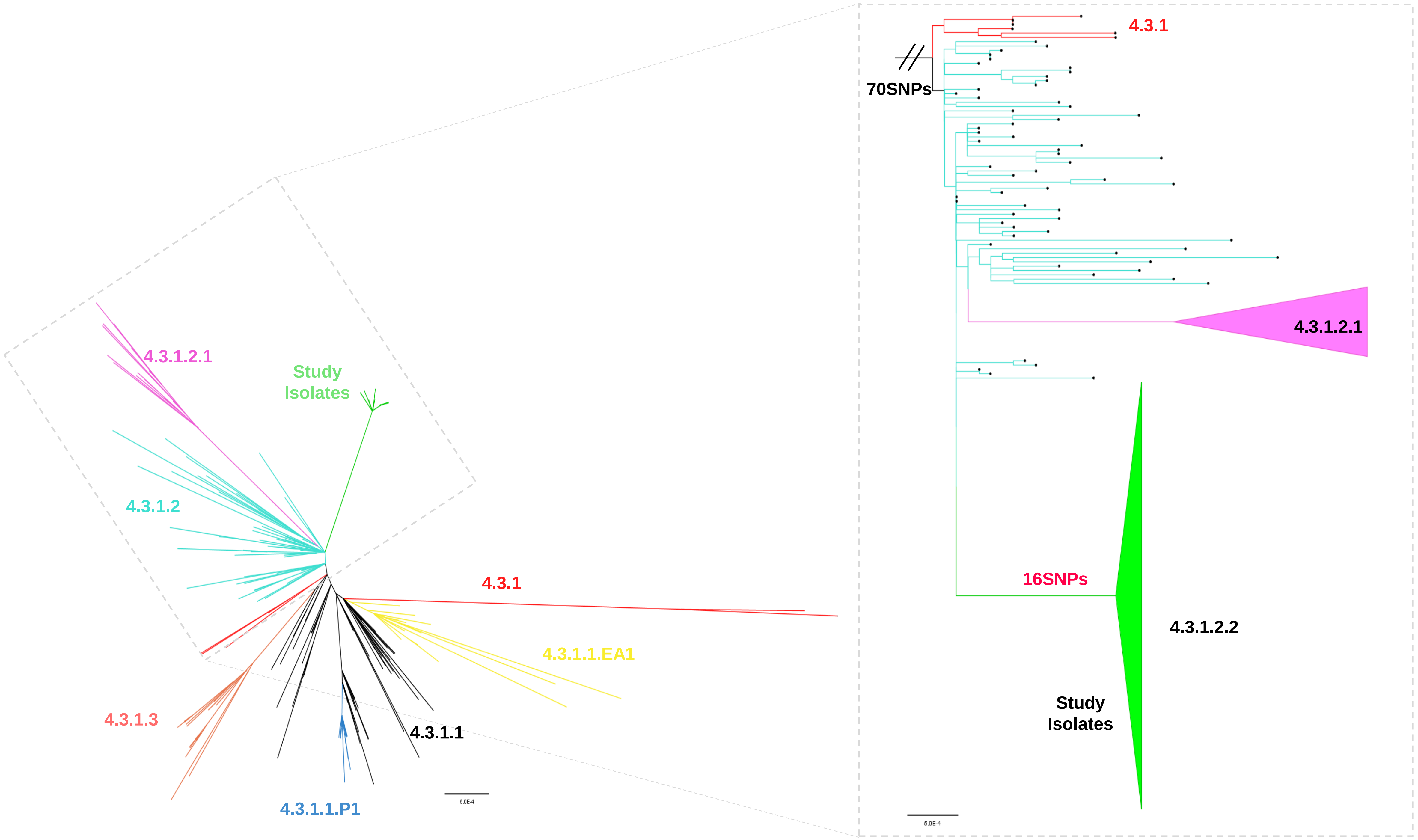
